# Effects of Oxygen Plasma Treatment on Parylene C and Parylene N Membrane Biocompatibility for Tissue Barrier Models

**DOI:** 10.1101/2022.06.09.495451

**Authors:** Shayan Gholizadeh, Daniela M. Lincoln, Zahra Allahyari, Louis P. Widom, Robert N. Carter, Thomas R. Gaborski

## Abstract

Porous membranes are integral components of in vitro tissue barrier and co-culture models and their interaction with cells and tissues directly affects the performance and credibility of these models. Plasma-treated Parylene C and Parylene N are two biocompatible Parylene variants with perceived potential for use in these models but their biocompatibility and biological interactions at their interface with cells are not well understood. Here, we use a simple approach for benchtop oxygen plasma treatment and investigate the changes in cell spreading and extracellular matrix deposition as well as the changes in material surface properties. Our results support the previous findings on the persistent effects of plasma treatment on Parylene biocompatibility while showing a more pronounced improvement for Parylene C over Parylene N. It is observed that although both increased surface roughness and persistent increases in oxygen species govern the plasma-driven improvement, the changes in oxygen concentration at the interface are the dominant factor. Overall, the results of this study provide a clear picture of potential mechanisms of plasma-induced changes in synthetic polymers which have implications for their use in in vitro model systems and other BioMEMS applications.

## 1. Introduction

There is an increasing interest in developing tissue-on-a-chip and barrier models, also known as microphysiological systems, due to their applications in drug discoveries and therapeutic strategies^1–3^. The blood-brain barrier (BBB) is highly important among barrier models^4^. In vivo, the BBB limits drug delivery to the brain, and its dysfunction has been shown to play a major role in the onset of neurodegenerative diseases^5,6^. Given the prevalence of neurodegenerative diseases such as Alzheimer’s disease, the necessity for effective brain drug delivery, and limited treatment options, platforms such as physiologically representative in vitro BBB models are highly sought for disease modeling and drug discoveries^7–9^. Porous membranes are integral components of these in vitro models^2,7,10–12^.

In barrier models, porous membranes serve as interfaces to establish a compartmentalized co-culture system between endothelial cells and another relevant cell type^13–19^. The two cell types are often grown on different sides of the membrane, and membrane pores facilitate paracrine signaling and direct physical contact^15,20^. Polymeric track-etched membranes are usually embedded in versatile culture inserts such as Transwell^®^ systems, and they are available with various pore sizes and porosities^2,15^. These are the most common membranes used for vascular barrier modeling^2^. However, these membranes have significant deficiencies, including limited live-imaging ability, irregular pore geometries, limited porosity, and, most importantly, thicknesses ranging from 5-10 μm (as opposed to the sub-micron endothelial-glial spacing observed in vivo) that significantly hinder cell-cell contact^15^. These shortcomings motivated the development of ultrathin nanomembrane technologies^2,15,21^.

Ultrathin nanomembranes often allow live imaging thanks to their optical transparency, allow effective cell-cell crosstalk due to thinness, and have higher porosities than their conventional membrane counterparts^15,20,21^. Despite these advances, almost all ultrathin and conventional membranes are made of synthetic materials with physical, chemical, and biological properties different from their biological analogs^2,12,14^. Remedies that compensate for this difference include membrane preincubation with cell culture media, extracellular matrix (ECM) coating, and plasma treatment^2,13,15,20,21^. Plasma treatment can be a more reproducible and effective method compared to other techniques to improve the biocompatibility of porous membranes^22–24^. However, it also encompasses complications such as transient effects and hydrophobic surface recovery and should be further investigated for different materials^15,25,26^.

It has been shown that plasma treatment affects organic and inorganic materials by enriching the surface with new oxygen-containing groups^27–30^. These effects fade relatively quickly, limiting the timeframe between membrane treatment and cell culture and creating inconsistent results due to time variability^31–33^. However, plasma treatment also creates surface roughness and micro- and nanotopographies as well as permanent changes in surface chemistry on organic materials such as synthetic polymers, which have been shown to improve cell attachment and biocompatibility through improving cell-substrate interactions^15,34,35^. Since well-attached cells deposit and assemble their own extracellular matrix (ECM) on the substrate, this can help mimic an in vivo microenvironment and be advantageous in creating a more physiologically representative barrier^7,36,37^. However, plasma treatment should be further explored and better understood for organic membrane material candidates in terms of physical and biological consequences. Parylene is among these candidate materials, and it has been previously examined for tissue barrier modeling^15,16,38^.

Parylene is a biocompatible synthetic polymer that has been approved by the Food and Drug Administration (FDA) for applications within the human body (USP XXII Class VI biocompatibility rating)^39,40^. Parylene has been successfully used for nanomembrane fabrication by our group as well as others, and it has generally shown favorable cell attachment and growth characteristics, especially after plasma treatment^15–17,41^. As part of our previous study, we briefly investigated whether the positive effects of plasma diminished over time by shelving the membranes for 7 or 14 days after treatment and before use. Interestingly, cell attachment remained high even after 14 days of storage^15^. However, a deeper understanding of the plasma effects is needed to achieve an ideal membrane for barrier models, and the effects of induced roughness and surface chemistry should be differentiated.

In this study, we investigated the effects of oxygen plasma treatment on two commonly used biocompatible Parylene variants, Parylene C and Parylene N, and compared their results with tissue culture polystyrene (TCPS) as well as silicon dioxide (SiO_2_) which is a commonly used inorganic material in BioMEMS applications. We switched from a reactive ion etching (RIE) tool used in our previous study to a more accessible benchtop plasma machine for facilitating future potential studies derived from this work. We assessed two endothelial cell types commonly used for barrier modeling to investigate the implications of varied plasma treatment lengths on cell-substrate interactions. We then characterized the surface in terms of roughness, contact angle, and oxygen species content. The results of this work can be expanded to any use of Parylene as a biocompatible material as well as other similar organic materials used as interfaces for cellular barrier and co-culture applications.

## 2. Methods

### 2.1 Membrane Preparation

Parylene membrane fabrication was conducted based on a protocol described previously^15^. Briefly, Micro-90, a water-soluble detergent, was used as a sacrificial layer deposited by spin-coating on 6-inch silicon (100) wafers. Parylene-C and Parylene N coating was done using DPX-C and DPX-N dimers (Specialty Coating Systems, USA), respectively, in an SCS Labcoter 2 Parylene deposition system (PDS 2010, Specialty Coating Systems, USA) (**Figure 1**)A. The process began at a base chamber pressure of 10 mTorr, and the dimer-cracking furnace was heated to 690 °C for Parylene C and 650 °C for Parylene N. Then, the vaporizer was ramped to a final temperature of 175 °C or 160 °C for Parylene C and Parylene N, respectively, causing the sublimation of dimer. The temperature ramp rate of the vaporizer was controlled to maintain a chamber pressure of 25 mTorr.

**Figure 1.**
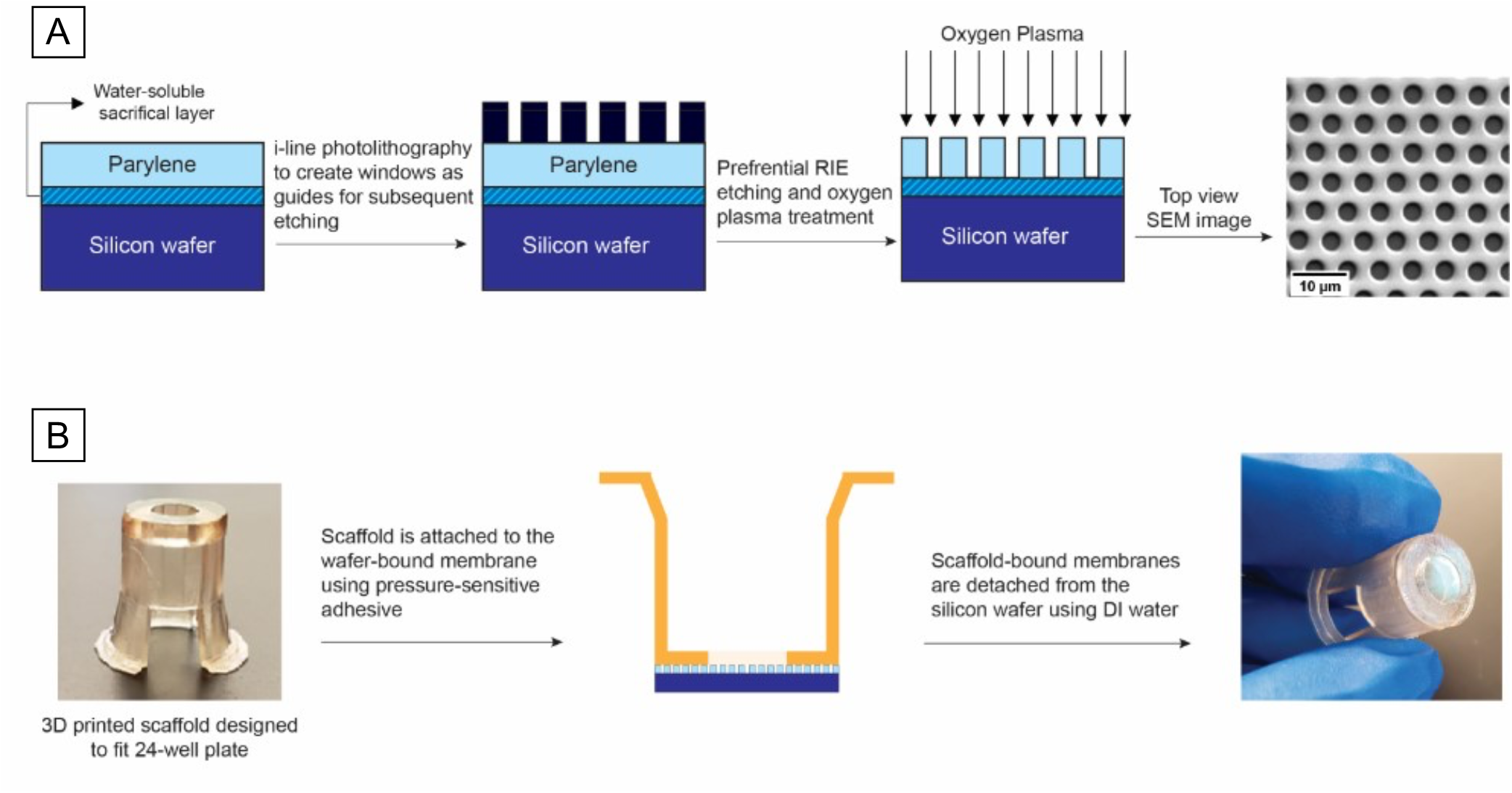
Fabrication, assembly, and plasma treatment of Parylene C and Parylene N nanomembranes for subsequent surface analysis and cell culture.

### 2.2 Oxygen Plasma Treatment

Unlike previous attempts to use high-cost sophisticated equipment with limited availability, such as inductively coupled plasma RIE tools, we aimed to use a benchtop plasma treatment tool that was more readily available. For this purpose, a Harrick PDC-001 plasma cleaner (Harrick, NY, USA) was used at its high-power setting for either 10 or 20 min, based on previous use of this tool for surface activation (**Figure 1B**). Samples were kept for 1-7 days for subsequent experiments before use.

### 2.3 3D Printing Scaffolds and Membrane Assembly

A Form 3B 3D printer (Formlabs, MA, USA) was used to fabricate cell culture scaffolds capable of supporting cell culture in multi-well plates. Briefly, a 3D mechanical design software (SolidWorks, MA, USA) was used to design scaffolds fitting in 24-well plates. The scaffolds were fabricated using BioMed Amber Resin (Formlabs, MA, USA) with resolution set to 50 μm. Smooth laser-cut polymethylmethacrylate (PMMA) circular parts were attached to double-sided pressure-sensitive adhesive (MP468, 3M, MN, USA) on both sides and were pushed on pieces of membrane-coated silicon wafers. Parylene membranes attached to the 3D-printed scaffolds were released by dipping in deionized (DI) water from the silicon substrate (**Figure 1B**).

### 2.4 Cell Culture

Pooled human umbilical vein endothelial cells (HUVECs) and a human cerebral microvascular endothelial cell line (hCMEC/D3) were cultured in EGM™-2 MV consisting of EBM™-2 and EGM™-2 MV SingleQuots™ Kit with 2.4% FBS, and 1% penicillin and streptomycin. HUVECs and cell culture reagents were purchased from Lonza (Walkersville, MD, United States). hCMEC/D3 cells were purchased from EMD Millipore (Temecula, CA, USA). HUVECs and hCMEC/D3 cells were detached by TrypLE and seeded on samples at a density of 2 × 10^4^ cells/cm^2^. Cells were used between passages 4-6.

### 2.5 Cell Spread Area

Cells were seeded on the Parylene C and Parylene N samples for 24 h, fixed in 3.7% formaldehyde for 15 min, and washed with PBS. This was followed by cell permeabilization for 3 min in 0.1% Triton X-100 and washing with double-distilled water. To visualize nuclei and cytoskeletons, HUVECs were stained using DAPI (300 nM) and 1:400 Alexa Fluor 488 conjugated phalloidin for 3 and 15 min, respectively. Cells were washed with PBS. Images were captured at 20× magnification through DAPI and GFP filters on a Keyence BZ-X700 microscope (Keyence Corp. of America, MA, USA). The area of spreading of actin fibers was measured for each cell using ImageJ software.

### 2.6 Fibronectin Fibrillogenesis

Fibronectin fibrillogenesis was evaluated on Parylene C and Parylene N samples. After cell culture for 24 h, cells were fixed with 3.7% formaldehyde for 15 min, washed with PBS, blocked with 20 mg/mL BSA for 15 min, and then washed again with PBS. Cells were stained using a 1:100 dilution of Alexa Fluor 488 conjugated anti-fibronectin (clone FN-3) for 2 hr, and washed with PBS three times. PBS was added to the samples, and they were imaged through a GFP filter using a Keyence BZ-X700 microscope. The lengths of fibrils were measured using the CT-FIRE package^42^, which was developed by the Laboratory for Optical and Computational Instrumentation at the University of Wisconsin-Madison to automatically extract individual collagen fibers from images.

### 2.7 Collagen IV Deposition

Collagen IV deposition was also evaluated on Parylene C and Parylene N samples through immunofluorescence microscopy. Similar to the fibronectin fibrillogenesis assay, cells were fixed with 3.7% formaldehyde for 15 min, washed with PBS, blocked with 20 mg/mL BSA for 15 min, and then washed again with PBS. Samples were incubated with 1:100 diluted Fluor 647 conjugated anticollagen IV (clone 1042) for 4 h and imaged through a Cy5 filter using a Keyence BZ-X700 microscope. Collagen IV coverage was quantified using the thresholding function of ImageJ software.

### 2.8 Contact Angle Measurement

The hydrophilicity of the surface of Parylene C-coated glass coverslips was evaluated through static contact angle measurements. For each condition, three different samples were used. 10 μl of DI water was placed at the center of the samples, and images were taken. Edges were omitted as these are typically also avoided in all microfabricated samples for reliability. The mean contact angle was extrapolated from three measurements for each sample via ImageJ (National Institutes of Health, USA).

### 2.9 Atomic Force Microscope Measurement

Parylene C and Parylene N membranes (untreated, treated for 10 min, and treated for 20 min) were examined using a MultiMode 8-HR atomic force microscope (AFM) (Bruker, USA). TR800PSA cantilevers (Oxford Instruments) with a spring constant of 0.15 N/m were utilized to scan the surface in tapping mode. The samples were prepared, mounted onto the AFMcell chamber, and surface topography scans were taken with a scan speed of 0.5 Hz to minimize imaging artifacts over a fixed area.

### 2.10 X-ray photoelectron spectroscopy (XPS)

X-ray photoelectron spectroscopy (XPS) was used to evaluate the effects of plasma treatment on the surface chemistry (1-10 nm depth) and its longevity over 7 days. XPS spectra were collected using a Kratos Axis Ultra DLD XPS system (Kratos Analytical, UK). After chamber pump-down and surface contaminant removal, survey scans with 1 eV steps and high-resolution scans for oxygen and carbon with 0.1 eV steps were collected. Shirley background was used to correct collected spectra and CasaXPS was utilized to deconvolute peaks and correct the spectra by shifting the carbon-carbon peak to 284.8 eV. The area under the curve for the oxygen peak was normalized to the corresponding area for aliphatic C and the oxygen peak area was used as a normalized representation of oxygen surface concentration.

## 3. Results and Discussion

### 3.1 Cell Spreading

Two endothelial cell types commonly used for vascular barrier studies (hCMEC/D3 and HUVEC) were used for cell-substrate interaction experiments. Instead of cell viability, we aimed to study cell spread area as a more direct measure of cell-substrate interaction. Our results demonstrated that plasma treatment increased the cell spread area for both treatment times, while more extended treatment led to marginally higher spread area for both hCMEC/D3 cells and HUVECs (**Figure 2**).

**Figure 2.**
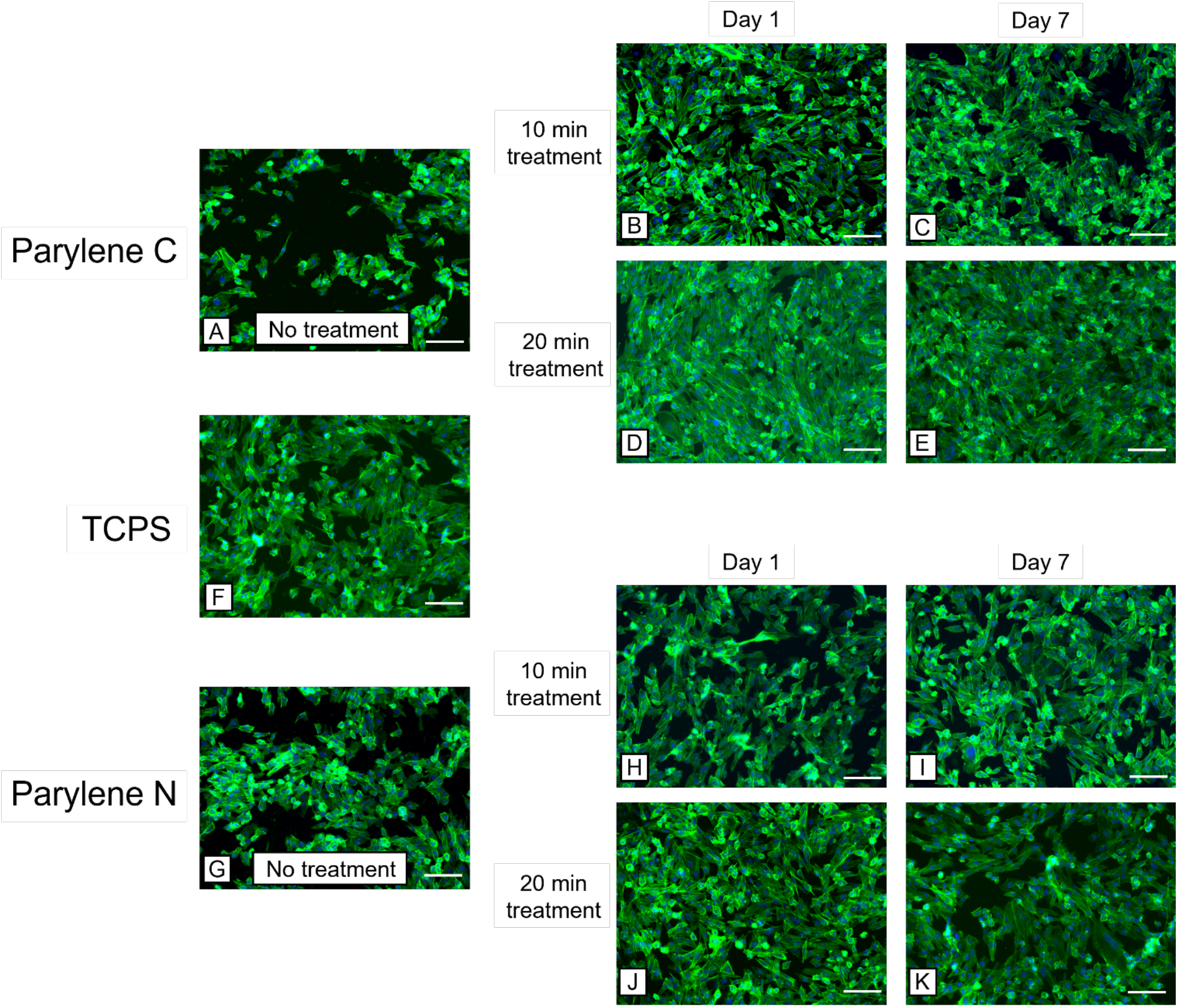
Representative images of nuclei (DAPI, blue) and F-actin (phalloidin, green) after 24 h on A-E) Parylene C membranes, F) TCPS, and G-K) Parylene N membranes. Scale bar = 100 μm in all images.

Figure 3 demonstrates cell spread analysis of HUVECs and hCMEC/D3 cells on Parylene C and Parylene N with different treatment and test conditions. It can be noted that although untreated Parylene C showed the smallest cell spread area, treated Parylene C membranes showed the highest cell spread area as compared to Parylene N membranes and TCPS. Another critical finding was that even though there was a slight effect of hydrophobic recovery (Figure 2C and 2K) in treated Parylene samples, there was no decrease in cell spread area on day 7 samples as compared to those samples seeded on day 1 after treatment, possibly due to permanent changes in surface properties.

**Figure 3.**
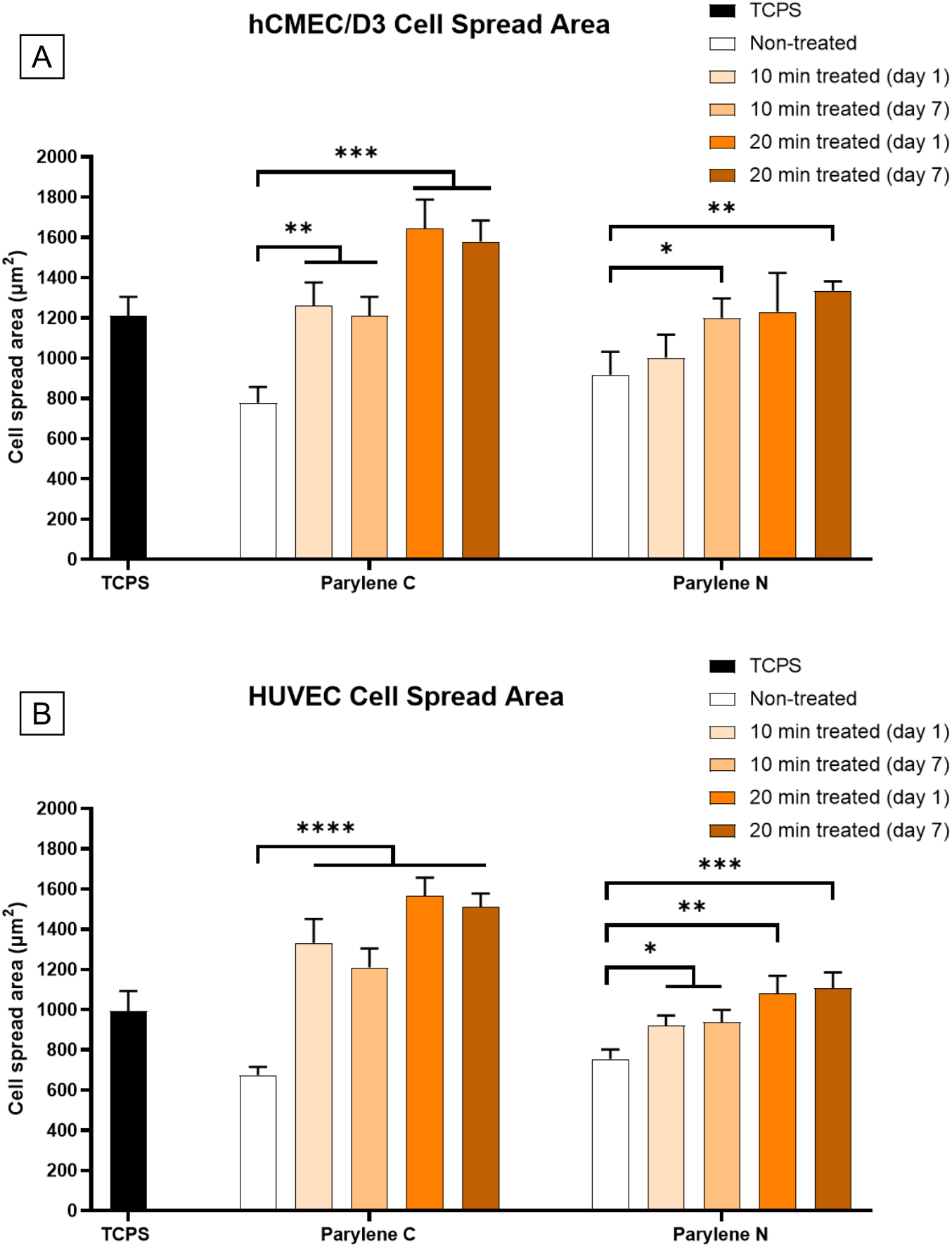
Changes in cell spread area on Parylene C and Parylene N membranes. A) hCMEC/D3 cell spread area on TCPS, Parylene C and Parylene N membranes. B) HUVEC cell spread area on TCPS, Parylene C and Parylene N membranes.

Another important observation is that there was no statistically significant improvement with 20 min treatment as compared to 10 min treatment except for hCMEC/D3s on Parylene C. This likely points to a saturation-like event either in surface roughness or surface bonds which will be explored in material characterizations in the following sections.

### 3.2 Fibronectin Fibrillogenesis

Fibronectin fibrillogenesis is associated with a cell’s ability to generate high traction forces^43–46^. Disruption in fibronectin fibrillogenesis can indicate lesser cell-substrate interaction and lower integrity in the formed endothelial barrier. Our results on fibronectin fibrillogenesis on hCMEC/D3 cells and HUVECs indicated that treatment of Parylene C and Parylene N led to a similar level of improvement in fibronectin fibrillogenesis (**Figures 4 & 5**).

**Figure 4.**
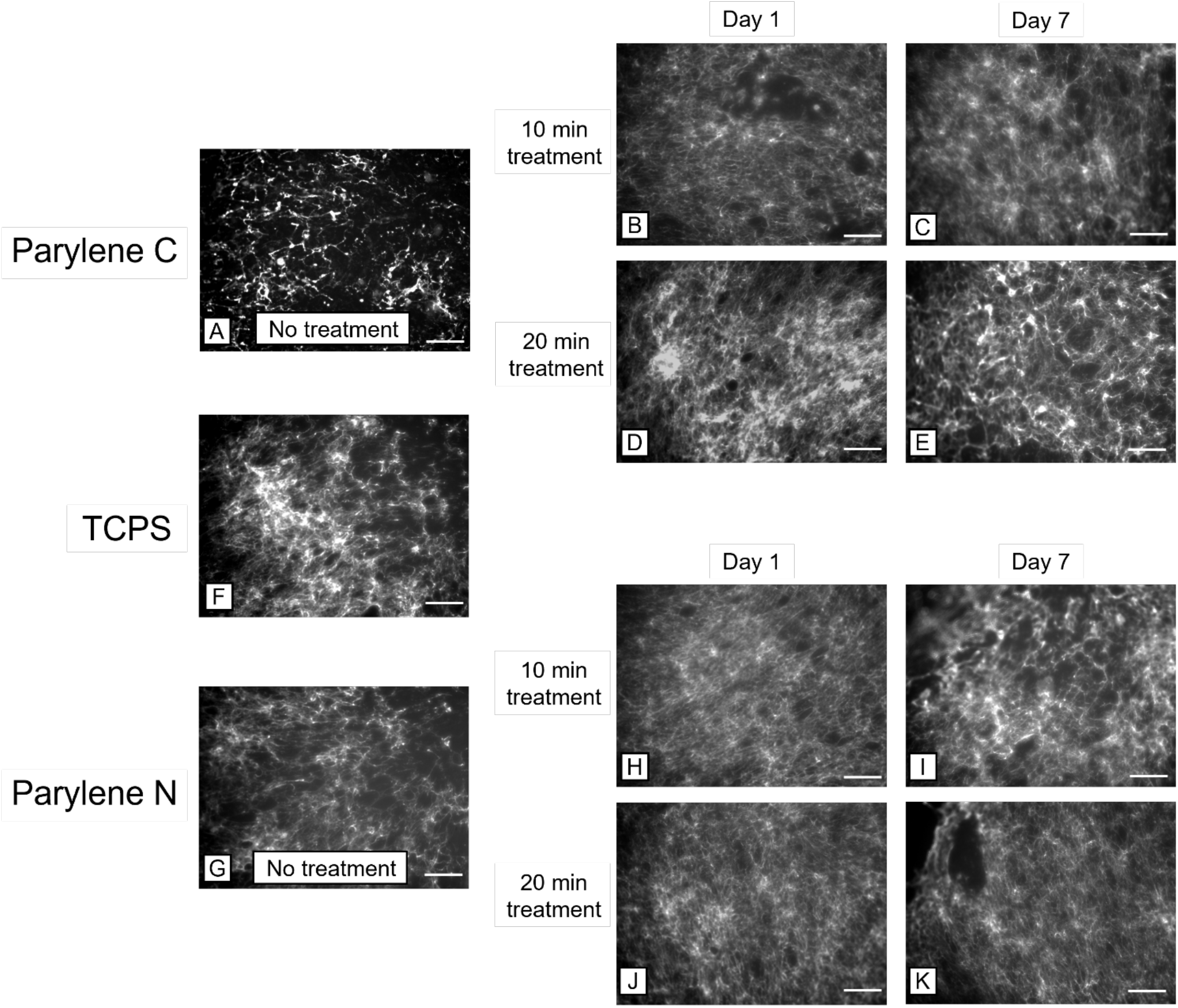
Representative images of fibronectin fibrillogenesis on A-E) Parylene C membranes, F) TCPS, and G-K) Parylene N membranes. Scale bar = 100 μm in all images.

**Figure 5.**
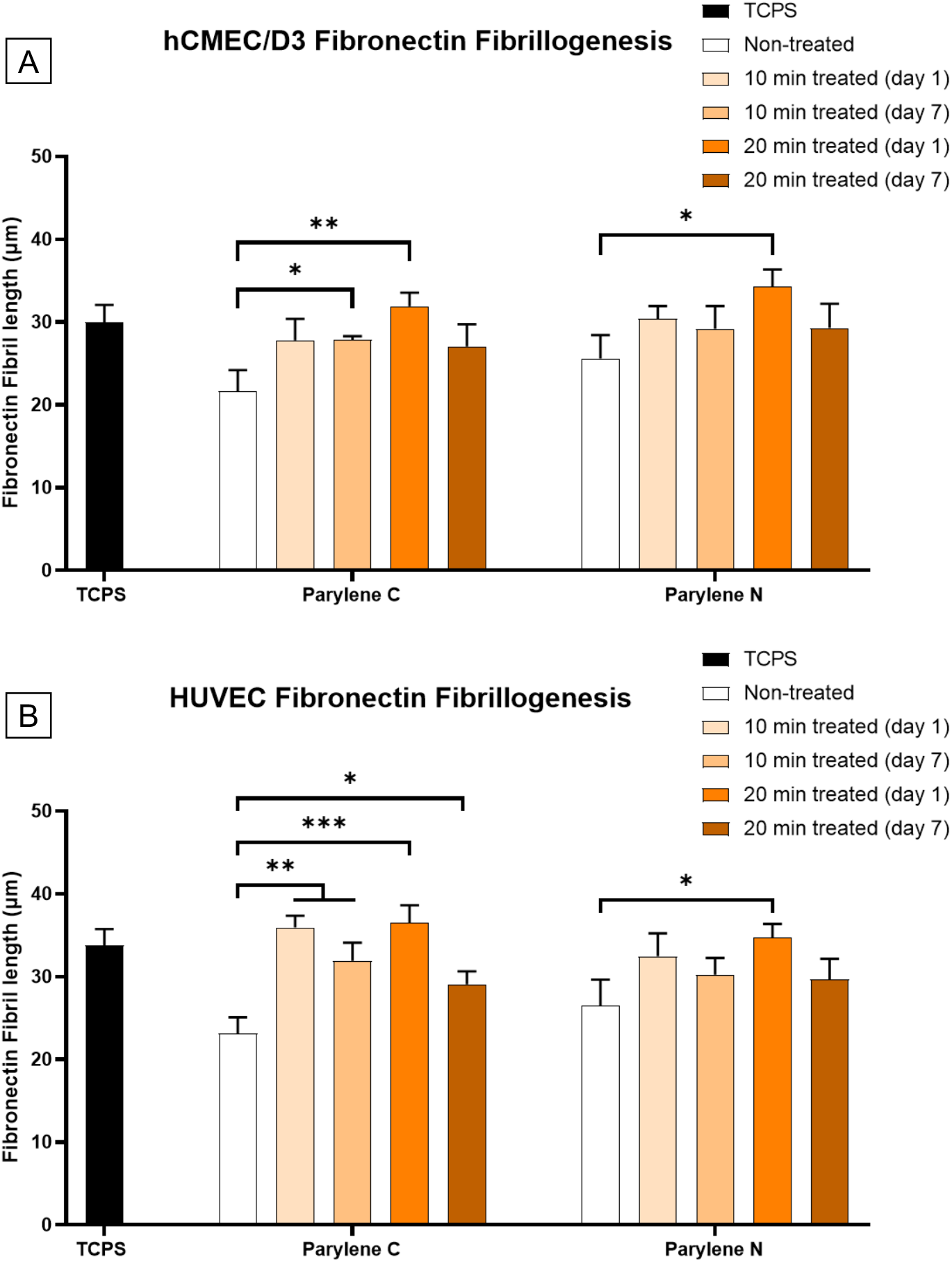
Changes in fibronectin fibrillogenesis on Parylene C and Parylene N membranes result from plasma treatment. A) hCMEC/D3 fibronectin fibril lengths on TPCS, Parylene C and Parylene N membranes. B) HUVEC fibronectin fibril lengths on TCPS, Parylene C and Parylene N membranes.

However, there is a noticeable decrease in fibronectin fibrillogenesis in most cases from day 1 to day 7, indicating that even with the perceived role of surface roughness, there is a somewhat apparent deterioration in the activated substrate. This decrease is only statistically significant in 20 min treated samples, indicating a baseline of surface activation that remained persistent. Despite the presence of this decrease, the results in all treatment conditions were still comparable to TCPS on day 7.

Another important observation is that only some of the day 1 samples were able to outperform TCPS in terms of fibrillogenesis. This contrasts cell spreading, where many of the treated Parylene samples demonstrated larger cell areas. All 16 sets showed statistically significant fibrillogenesis enhancement over untreated samples, cementing the persistent nature of Parylene plasma activation.

### 3.3 Collagen IV Protein Expression

The interaction of endothelial β1-integrins with collagen IV of the basement membrane is correlated to the expression of the tight junction protein claudin-5 and barrier integrity in vitro^47^. The interaction through the integrin receptors provides physical support and regulates signaling pathways, whereby the endothelial cells can adapt to changes in the microenvironment. Therefore, the ability of endothelial cells to deposit collagen IV can be vital to having more physiologically representative in vitro barrier models.

Our results display significant improvement in collagen IV deposition due to plasma treatment across all samples, and they either perform comparably or better than TCPS (**Figures 6 & 7**). The decrease from day 1 to day 7 is still observed in 20 min treated samples but is much less pronounced than for fibronectin fibrillogenesis. The results are promising for in vitro barrier models in terms of reliability, persistency, and effectiveness of plasma treatment for collagen IV deposition.

**Figure 6.**
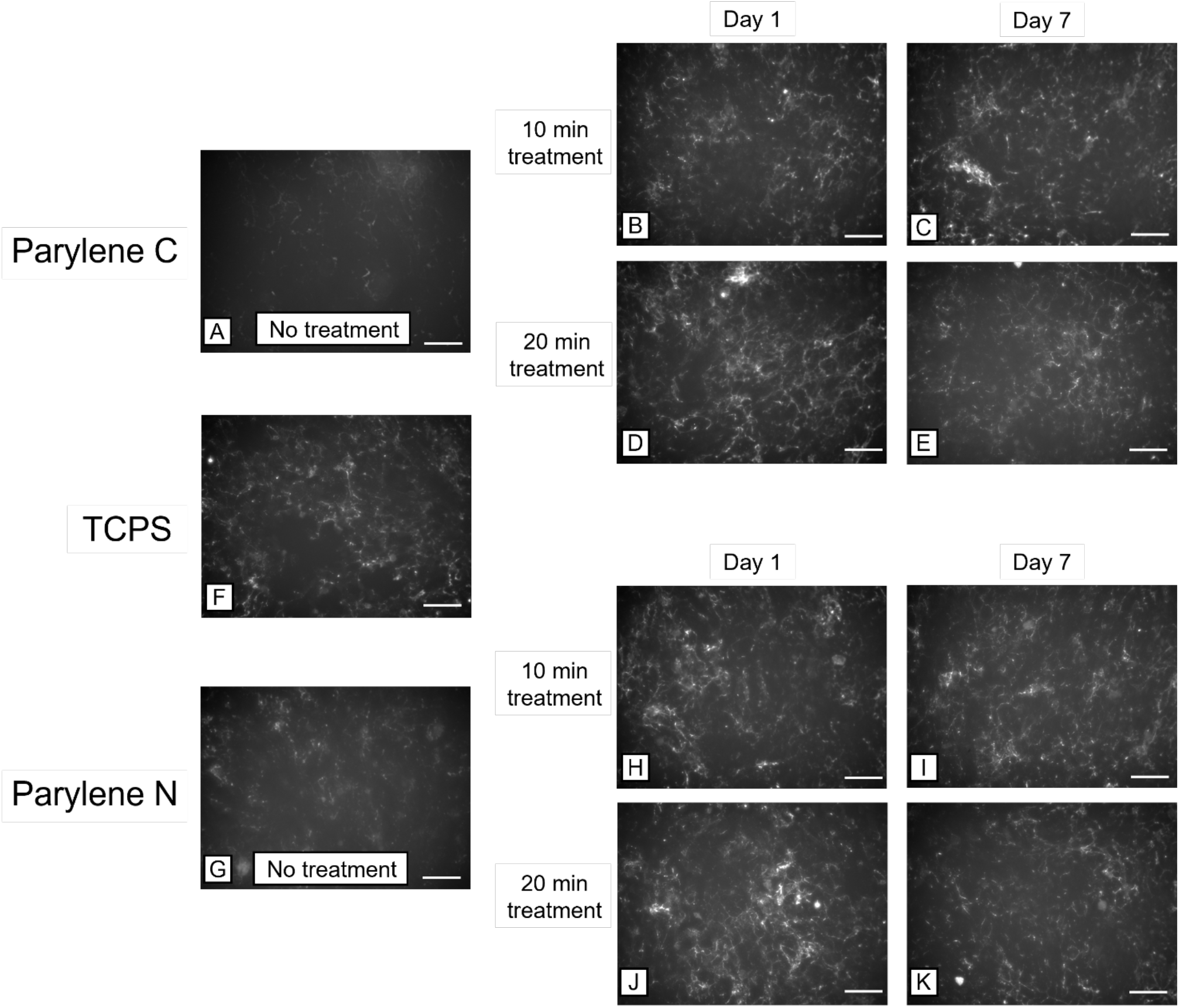
Representative images of Collagen IV deposition on A-E) Parylene C membranes, F) TCPS, and G-K) Parylene N membranes. Scale bar = 100 μm in all images.

**Figure 7.**
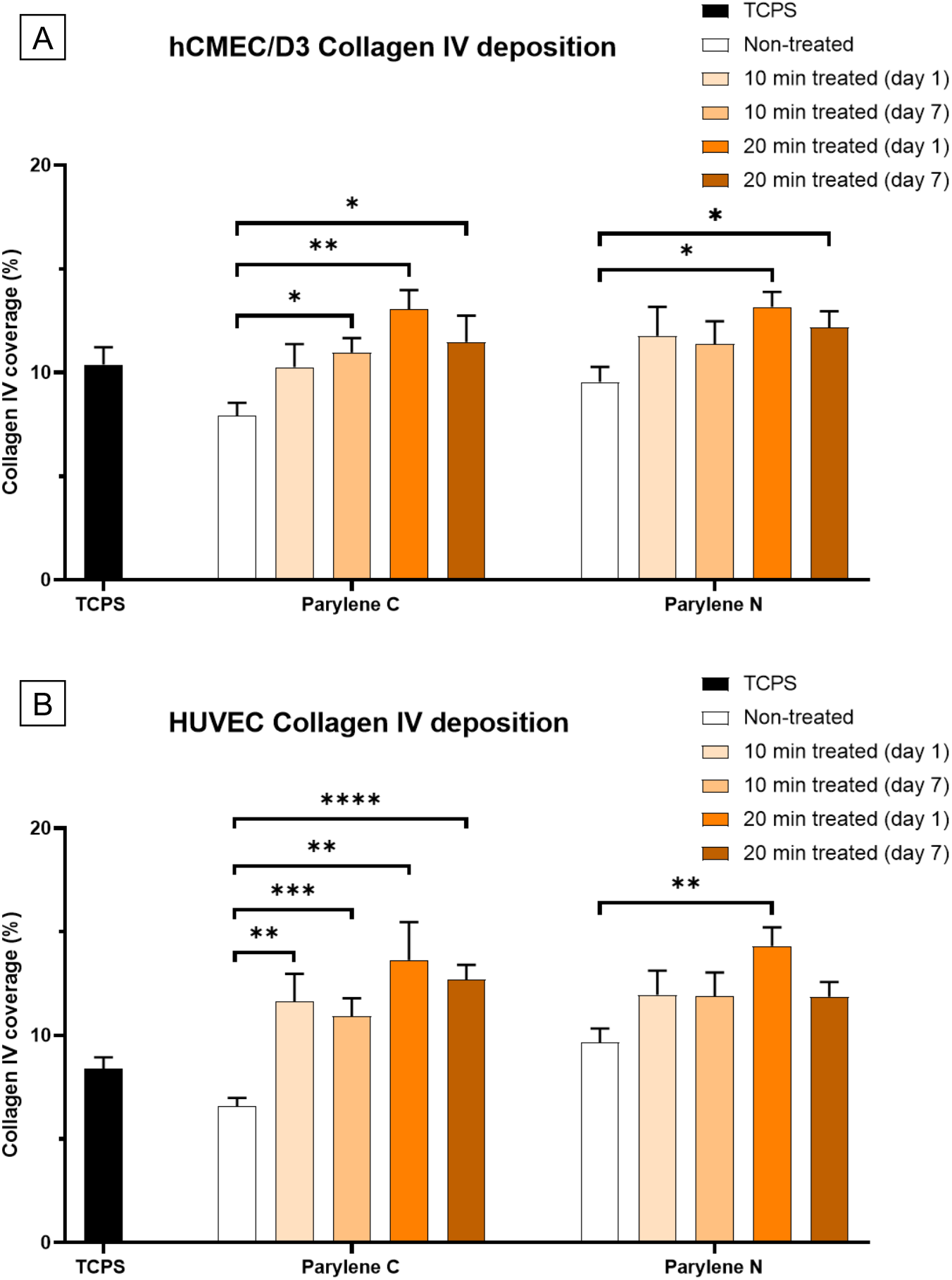
Changes in collagen IV deposition on Parylene C and Parylene N membranes result from plasma treatment. A) hCMEC/D3 collagen IV coverage on TCPS, Parylene C and Parylene N membranes. B) HUVEC collagen IV coverage on TCPS, Parylene C and Parylene N membranes

### 3.4 Hydrophilicity and Hydrophobic Recovery

It is well-known that hydrophilic surfaces promote cell adhesion. Given the prevalent use of inorganic surfaces such as silicon dioxide (SiO_2_) for cell culture and the smooth nature of these materials, they are good candidates for comparison to see the extent of time-persistent surface plasma effects. Day 1, day 3, and day 7 measurements following plasma treatment show that Parylene C and Parylene N both exhibit much lower contact angles and higher hydrophilicity compared to untreated samples, and they retained this characteristic with very slight hydrophobic recovery (**Figure 8**). However, smooth SiO_2_ substrates showed almost complete recovery by day 7, indicating the possible role of roughness in retaining active sites and increasing hydrophilicity. A notable finding is that nearly all of the hydrophobic recovery across the three groups occurred within 3 days, and the decrease in hydrophobicity is minor after day 3.

**Figure 8.**
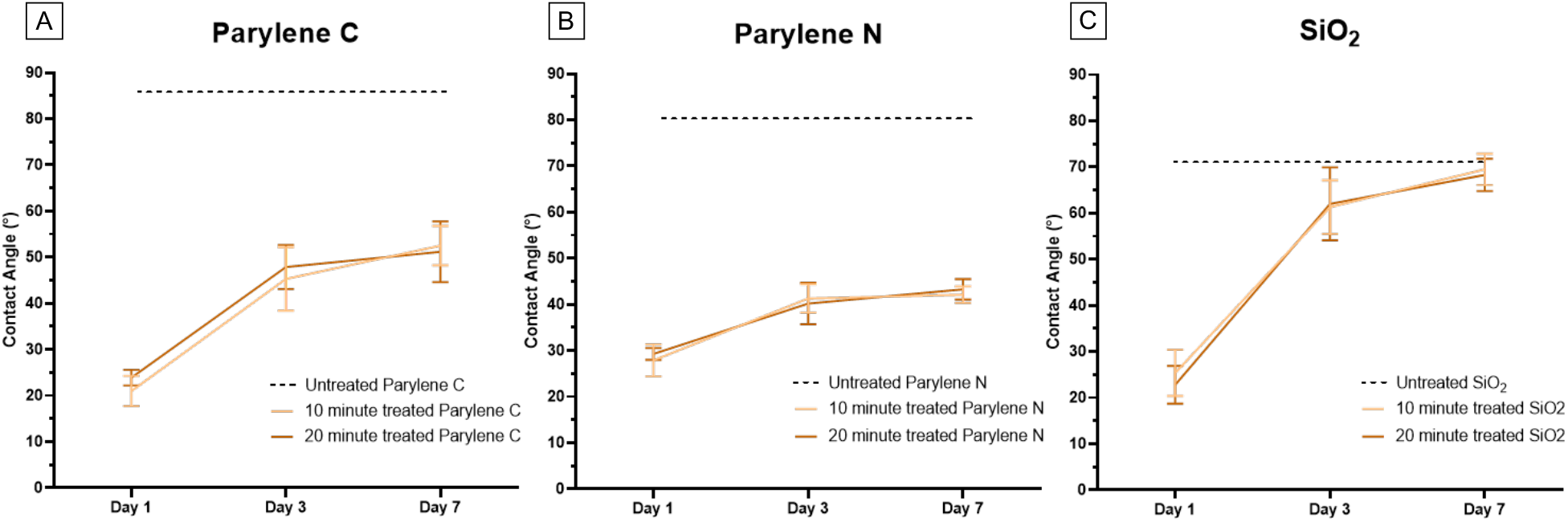
Contact angle measurements from day 1, day 3, and day 7 of plasma treatment for A) Parylene C, B) Parylene N, and C) SiO2 surfaces. The dashed line in each plot represents the measurements from the respective untreated material.

### 3.5 Surface Morphological Evaluation

Previous work on the underlying mechanisms of plasma-induced surface activation has mainly focused on surface energy and surface chemistry with valuable insights and outcomes^48,49^. However, the potential role of surface morphological changes has not received significant attention. In previous published works, it was observed that plasma activation had shown persistency over time to various extents, at least for organic surfaces^15,50,51^. Due to the often time-dependent nature of surface energy and chemistry changes, especially when exposed to air, said persistency was not fully explained.

Parylene C and Parylene N have minor structural differences, but AFM measurements demonstrate distinct differences in surface roughness between the two variants (**Figure 9**). Parylene C, as expected based on previous work, shows a slight increase in roughness as treatment time increased. Although this can partially explain the observed persistency for Parylene C membranes, such an increase in surface roughness was not observed for Parylene N. However, since all AFM visualizations were plotted on the same scale, it can be appreciated that untreated Parylene N exhibits higher surface area and rough sites than untreated Parylene C.

**Figure 9.**
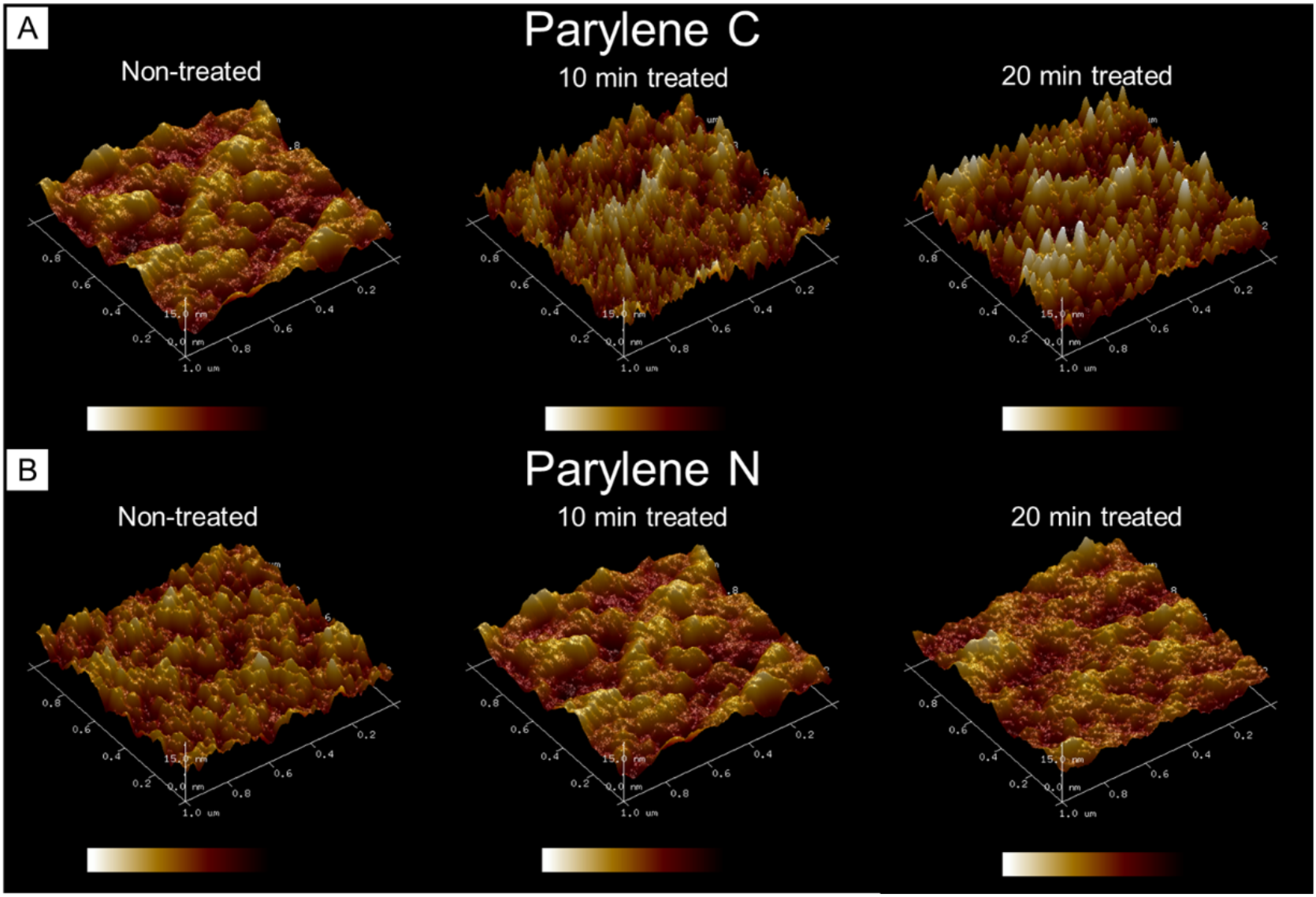
The effect of surface plasma modification of different lengths of time on surface topography and roughness. AFM images of parylene C and parylene N surfaces demonstrated slight increases in introduced surface area and roughness on plasma-treated parylene C as the treatment time increased. Noticeable changes were not observed in plasma-treated parylene N.

### 3.6 X-ray photoelectron spectroscopy (XPS)

As opposed to the surface topography analysis with AFM where only mild increases in Parylene C roughness were observed, XPS spectra analysis exhibited strong agreement with the observed improvement in the cell interactions with plasma-treated Parylene C and N membranes in the previous sections (**Figure 10**). In previous work, a correlation between the presence of oxygen-containing surface groups and improved cell biocompatibility was established^29^. Plasma treatment led to an increase in the surface oxygen bonds (C-O and C=O), while the 20 min treatment led to a slightly higher amount of oxygen species. This mirrored the observed changes in cell interactions on the treated Parylene substrates. 7 days after the oxygen treatment, plasma-induced improvement remained almost consistent across all of the samples. Similar to the more significant improvement in Parylene C interaction with endothelial cells as compared to Parylene N, the increase in oxygen species in Parylene C was more noticeable. This showed that changes in surface chemistry govern the cell-substrate interactions on Parylene C and Parylene N substrates and play a more significant role than surface topography, at least for endothelial cells typically used for barrier studies.

**Figure 10.**
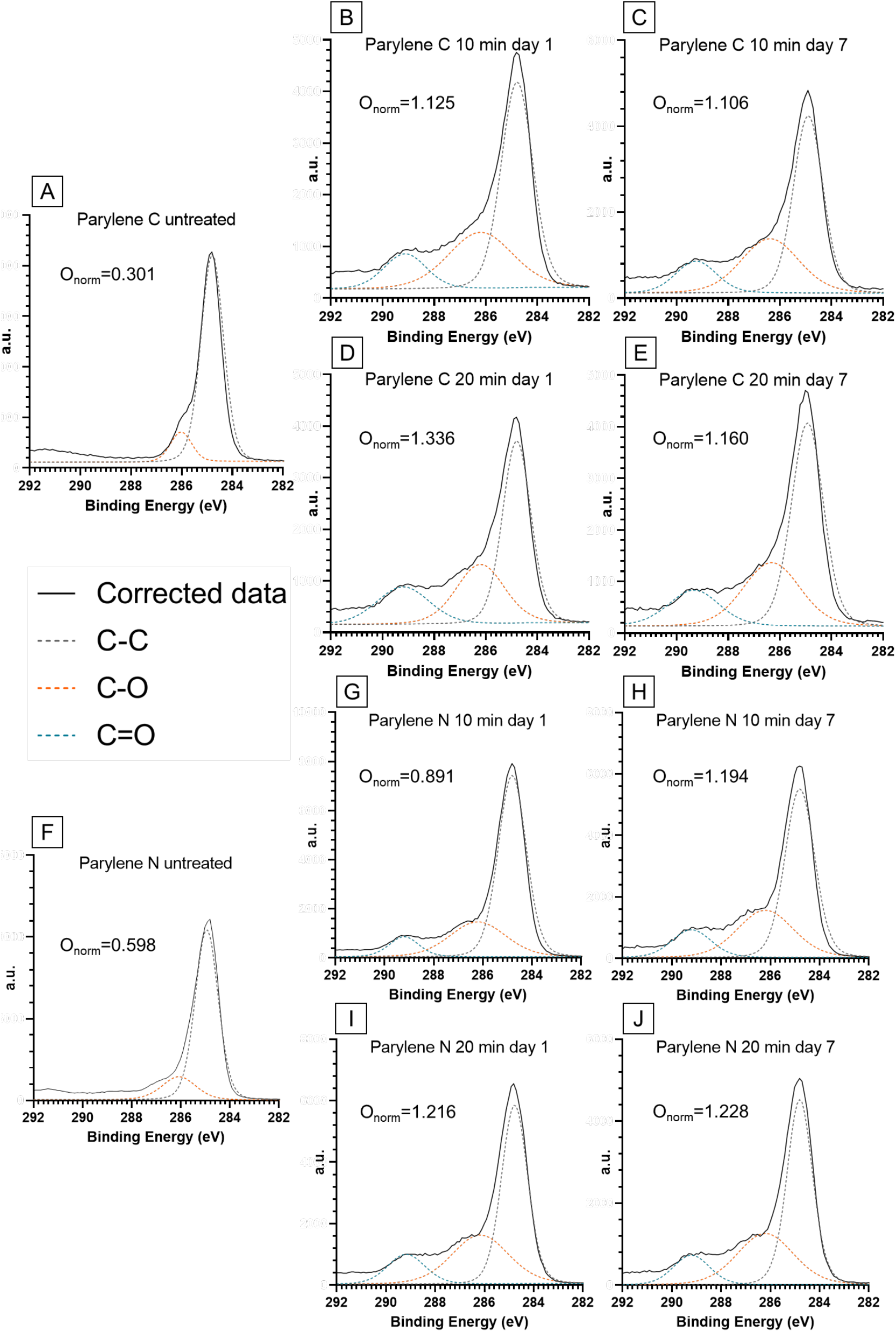
XPS visualization of high-energy tails associated with carbon bonding to oxygen for Parylene C (A-E) and parylene N (F-J) membranes with different treatments on different days and corresponding normalized oxygen concentration extracted from the survey scans (normalized to carbon at%). Continuous black lines in each graph correspond to the corrected data and the dotted lines are Gaussian-Lorentzian fits for carbon-oxygen (orange and blue) and carbon-carbon (gray) bonds.

## 4 Conclusion

We explored the time-dependent effects of oxygen plasma treatment, a less studied but important factor governing surface plasma activation. Oxygen plasma treatment is often a simple and effective method for increasing cell biocompatibility and surface activation. We conclusively showed that oxygen plasma treatment has a wide range of effectiveness for different formulations of Parylene. Our results and comparisons to smooth SiO_2_ substrates indicate that surface roughness may play a role in the effectiveness and longevity of oxygen plasma treatment. We demonstrated that using a readily available benchtop oxygen plasma cleaner can suffice for treating polymers such as Parylene C and N compared to more expensive techniques such as reactive ion etching. It was also shown that upon oxygen plasma treatment, hydrophobic recovery of a smooth surface such as SiO_2_ sharply contrast those of rough Parylene surfaces, indicating an essential consideration for using different material types for in vitro human models such as the BBB. It was observed that the improvement in cell-Parylene substrate interaction as a result of plasma treatment was mainly governed by the permanent surface chemistry changes while surface roughness likely plays a secondary and less significant role in these interactions. Future work should focus on functional studies of in vitro barrier models to directly investigate the role of surface treatment on endothelial barrier properties such as tight junction protein expression, small molecule permeability, and potential upregulation in the expression of BBB-specific genes.

## ACKNOWLEDGEMENTS

Research reported in this publication was supported by NIGMS of the National Institutes of Health under award number R35GM119623 to TRG. The content is solely the authors’ responsibility and does not necessarily represent the official views of the National Institutes of Health. We thank Daniel DiMartino for his assistance in cell culture.

